# Microtubule end tethering of a processive Kinesin-8 motor Kif18b is required for spindle positioning

**DOI:** 10.1101/286658

**Authors:** Toni McHugh, Agata Gluszek-Kustusz, Julie P.I. Welburn

## Abstract

Mitotic spindle positioning specifies the plane of cell division during anaphase. Spindle orientation and positioning is therefore critical to ensure symmetric division in mitosis and asymmetric division during development. The control of astral microtubule length plays an essential role in positioning the spindle. Here we show using gene knockout that the Kinesin-8 Kif18b controls microtubule length to center the mitotic spindle at metaphase. Using an integrated approach, we reveal that Kif18b is a highly processive plus end-directed motor that uses a C-terminal non-motor microtubule-binding region to accumulate at growing microtubule plus ends. This region is regulated by phosphorylation to spatially control Kif18b accumulation at plus ends and is essential for Kif18b-dependent spindle positioning and regulation of microtubule length. Finally, we demonstrate that Kif18b shortens microtubules by increasing the catastrophe rate of dynamic microtubules. Overall, our work reveals that Kif18b utilizes its motile properties to reach microtubule ends where it regulates astral microtubule length to ensure spindle centering.

## Introduction

Spindle positioning and orientation is essential to ensure accurate chromosome partitioning and symmetrical cell division. Proper spindle placement is also particularly important during development and in stem-cell homeostasis, when cells divide asymmetrically to specify cell differentiation and generate daughter cells of different cell size and fate (Siller and Doe, 2009). The length and density of astral microtubules influence the position of the spindle by altering the interactions between astral microtubules and cortical force-generators (Kiyomitsu and Cheeseman, 2012; Samora et al., 2011; Garzon-Coral et al., 2016). At the interphase to mitosis transition the microtubule cytoskeleton undergoes rapid remodeling. The increased dynamicity of microtubules allows the depolymerization of long interphase microtubules and subsequent assembly of dynamic spindle and astral microtubules that build and position the bipolar spindle (Belmont et al., 1990; Rusan et al., 2001).

Kinesin-8 and Kinesin-13 motors regulate microtubule dynamics and length across eukaryotes. However, the microtubule depolymerization mechanism of Kinesin-8 motors appears to differ across species. In budding yeast, Kip3 walks along microtubules and depolymerizes them (Gupta et al., 2006; Su et al., 2011; Varga et al., 2009), whereas Drosophila Klp67A localizes to kinetochores where it regulates spindle length (Savoian and Glover, 2010). Whether human Kinesin-8 Kif18a motor is a depolymerizing enzyme, a processive motor that dampens microtubule plus end dynamics, or both remains under debate (Mayr et al., 2007; Stumpff et al., 2008). A second human Kinesin-8, Kif18b, is reported to exhibit diffusion on the microtubule lattice using its C terminus and weak directed motility, which does not explain how it could target to or destabilise microtubule tips (Shin et al., 2015). Kif18b has previously been implicated in the negative regulation of astral microtubule length and has a modest contribution to chromosome alignment (Stout et al., 2011; Tanenbaum et al., 2011; Walczak et al., 2016). Kif18b requires EB1 for microtubule end accumulation but the EB-binding motifs in Kif18b are not sufficient for plus tip localization (Tanenbaum et al., 2011). Additionally, Kif18b may precede EB1 at microtubule ends (Shin et al., 2015), suggesting other mechanisms enable Kif18b targeting to microtubule ends. Whether Kif18b cooperates with the Kinesin-13 microtubule depolymerase MCAK or independently depolymerizes microtubule ends also remains under debate (Tanenbaum et al., 2011; Walczak et al., 2016).

Here we combine cell biology, biochemistry, and single molecule reconstitution assays to define the molecular mechanisms that allow Kif18b to differentially target and accumulate at microtubule ends, where it plays an important role in regulating microtubule length and spindle positioning. We demonstrate that Kif18b tracks the growing ends of microtubules autonomously *in vitro* and reduces microtubule length by promoting microtubule catastrophe. We propose that Kif18b uses its motile properties to reach and accumulate at microtubule ends in a phospho-specific manner to selectively destabilize astral microtubules.

## Results

### Kif18b and MCAK are major mitotic motors negatively regulating microtubule length

Microtubule length regulation plays an important role in spindle assembly, geometry and positioning. Previous work has analyzed the consequences of depleting kinesins that regulate microtubule length in human cells, but with differing results, possibly due to variable efficiencies of protein depletion or off-target effects (Bakhoum et al., 2009; Manning et al., 2007; Mayr et al., 2007; Tanenbaum et al., 2009; Welburn and Cheeseman, 2012). To identify kinesins that regulate microtubule length, we measured microtubule length in cells depleted for the Kinesin-13 members Kif2a, Kif2b, and Kif2c/MCAK and the Kinesin-8 members Kif18a and Kif18b using siRNA after Eg5 inhibitor treatment (Fig S1A, B). We found that both MCAK and Kif18b regulate microtubule length in mitotic cells, in agreement with previous reports (Fig S1C, D) (Stout et al., 2011; Tanenbaum et al., 2011; Walczak et al., 2016). However, Kif2a, Kif2b and Kif18a depletion did not alter microtubule length. In addition, co-depletion of Kif18b and MCAK did not have an additive effect on microtubule length, suggesting they work in the same pathway to regulate astral microtubules (Fig S1C, D).

To define the effect of Kif18b in regulating microtubule length, we generated a stable HeLa cell line lacking Kif18b using CRISPR/Cas9-mediated gene targeting, indicating that Kif18b is not essential for viability of cultured HeLa cells (see Methods). We found that Kif18b expression was eliminated from HeLa cells targeted by Cas9, while there was residual Kif18b protein upon RNAi depletion, which could mask the phenotype (Fig 1A). We then observed the mitotic microtubule organization of the cell line lacking Kif18b. Similarly to cells depleted for Kif18b using RNAi, we observed a strong increase in the length of astral microtubules (Fig 1B)(Stout et al., 2011; Tanenbaum et al., 2011; Walczak et al., 2016). This was particularly apparent when Aurora kinases A and B were simultaneously inhibited. In the absence of Aurora kinase activity, microtubules in HeLa cells were almost completely depolymerized, while they remain over 6 μm in the cell line lacking Kif18b (Fig 1C, D). This is consistent with the Aurora kinases negatively regulating Kif18b mediated microtubule destabilization (Tanenbaum et al., 2011).

**Figure 1:**
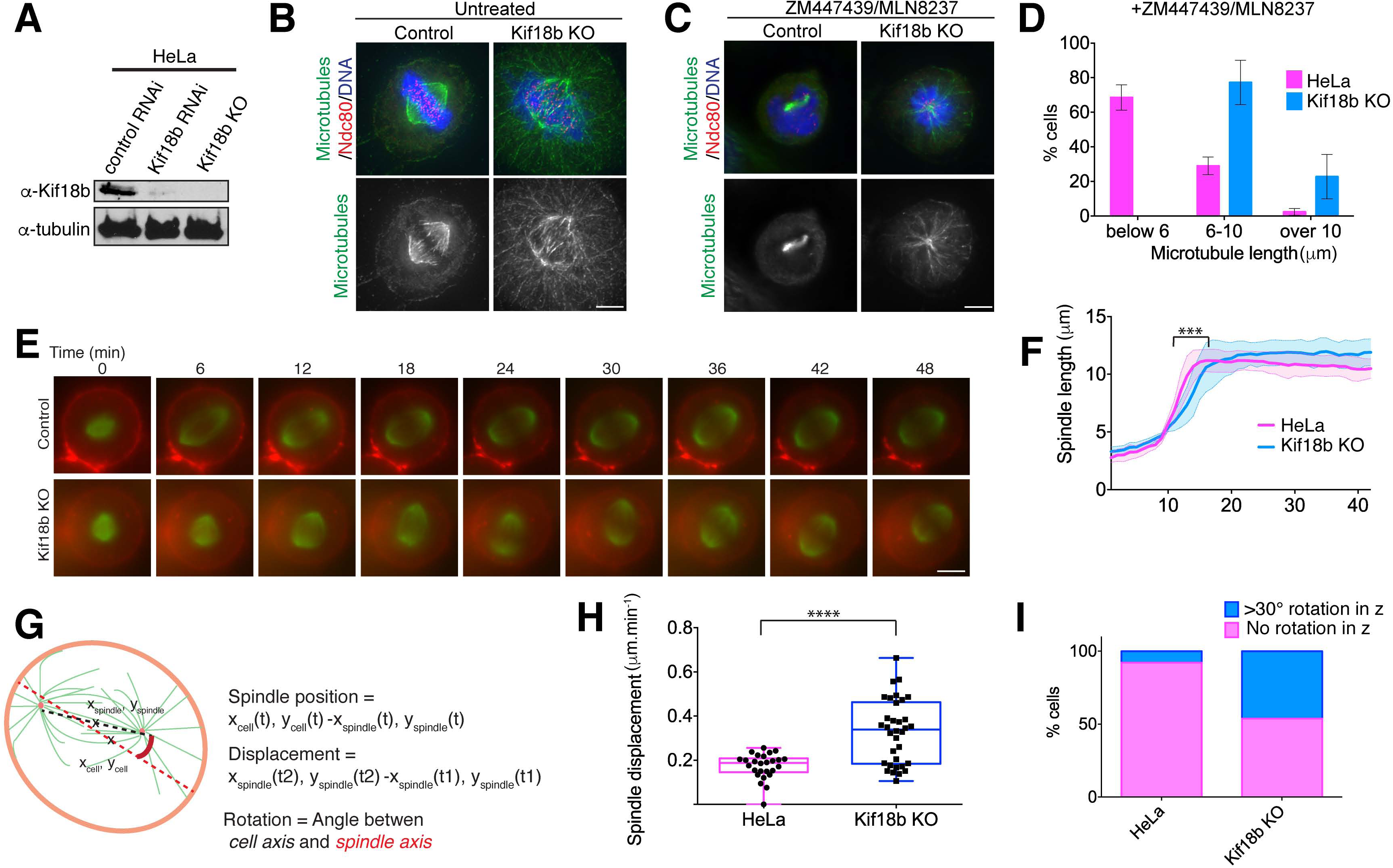
Kif18b controls astral microtubule length and spindle positioning. (A) Western blot showing depletion and gene knockout of Kif18b in HeLa cells using siRNA and CRISPR-Cas9. (B) Representative immunofluorescence images of control and Kif18b-KO HeLa cells, acquired using antibodies against Ndc80 and β-tubulin. Gamma level for the microtubule channel was altered to observe the astral microtubules. Images are scaled similarly. (C) Representative immunofluorescence images of control and Kif18b-KO HeLa cells treated with Aurora kinase A and B inhibitors MLN8237 and ZM447439 acquired using antibodies against Ndc80 and β-tubulin. (D) Quantification of microtubule length was measured in >30 cells per experiment. (E) Representative time-lapse imaging of cells incubated with SiR-tubulin and CellMask dye after an STLC washout and MG132 treatment. (F) Graph representing the average spindle length and the corresponding SD for HeLa and Kif18b-KO HeLa cells (n= 32 and 36), defined in (E) during spindle elongation. Delay in spindle elongation of Kif18b-KO cells was statistically significant. (G) Schematic diagram showing how spindle position, displacement and rotation are determined. (H) Box and whisker plot showing quantification of the displacement of the spindle from the center of the cell during metaphase. Each point represents the displacement of 1 spindle over at least 30 minutes (HeLa n=26 cells, Kif18b KO=34 cells). All error bars represent standard deviation. Asterix indicate Unpaired T-test test significance value. ***P<0.001, ****P<0.0001. Scale bars 5 μm. (I) Quantification of the number of cells undergoing more than 30**°** rotation in z out of the imaging plane during metaphase. (HeLa n=86 cells, Kif18b KO=122 cells).

### Kif18b ensures correct spindle positioning

To define the role of Kif18b-regulated astral microtubule length control on spindle assembly and positioning, we imaged mitotic cells using a previously described assay to examine spindle bipolarity establishment (Young et al., 2014). Overall, spindle elongation dynamics and length were similar in control cells and Kif18b knockout (KO) cells, although we observed a significant delay in reaching the maximum spindle length when Kif18b was absent (Fig 1E, F). The spindles of HeLa cells became bipolar and remained centered and parallel to the imaging plane, with a spindle displacement rate of 0.19 μm/min (±0.13 μm) (Fig 1G, H). In contrast, we observed frequent spindle rotation outside of the imaging plane (greater than 30**°**) in cells lacking Kif18b (Fig 1I). Kif18b KO spindles displayed strong positioning defects, with spindles moving around the cell centre and proximally to the cell cortex (movie 1). Spindles had a random oscillatory-like trajectory moving on average at 0.50 μm/min (±0.18 μm) within the cell (Fig 1G-I, Fig S1E, F). These results indicate that increasingly long astral microtubules cause spindle mispositioning and rocking, likely through additional interactions with force-generators at the cell cortex. At anaphase entry, the spindles of Kif18b KO cells were less centered than in control HeLa cells. During anaphase, the rate of spindle elongation and final spindle length for Kif18b KO cells were greater than for control HeLa cells, possibly because microtubules were longer (Fig S2A, B). In both Kif18b KO cells and control HeLa cells, the displacement of the spindle with respect to the centre of the cell decreased, resulting in the correction of spindle mispositioning (Fig S2A, C). Previous work has shown that an anaphase-specific dynein-dependent pathway centers the spindle as cells progress to anaphase (Kiyomitsu and Cheeseman, 2013). It is possible that the anaphase spindle positioning mechanism becomes essential when the length of microtubules increases, causing spindle mispositioning during metaphase. Taken together, our data indicate that Kif18b plays a role in reducing the length of astral microtubules to facilitate the correct positioning of the metaphase spindle.

### Kif18b has a C-terminal microtubule-binding domain, whose affinity is controlled by phosphorylation

Kif18b localizes to microtubule ends (Tanenbaum et al., 2011; Walczak et al., 2016). Although Kif18b requires EB1 to localize to microtubule ends, additional factors play a role in Kif18b plus end targeting (Stout et al., 2011; Tanenbaum et al., 2011). To define the unique targeting properties of Kif18b, we generated multiple Kif18b constructs (Fig 2A). First we observed that the GFP N-terminal tagging of Kif18b led to chromosome targeting of Kif18b and prevented its physiological localization to astral microtubules Fig S1E). This indicates that GFP at the N terminus of Kif18b may interfere with its motor properties. We then generated minimal Kif18b motor constructs lacking its C terminus with a C-terminal GFP. Kif18b-GFP displayed robust localization to microtubule plus ends (Fig 2B). In the absence of residues 591-852, the Kif18b motor only targets to spindle microtubules (Fig 2B). The kif18b motor only containing the dimerization domain targeted to spindle microtubules (Fig 2B). To test whether the C terminus of Kif18b was the main determinant in motor targeting, we swapped the tail domains of Kif18a and Kif18b. Despite highly conserved motor domains target to distinct cellular localizations, Kif18a targets to K-fibers and kinetochores, while Kif18b targets to microtubule plus ends (Fig 2B, C). We generated a chimeric kinesin consisting of the motor and neck-linker region of Kif18a fused to the C terminus of Kif18b. This Kif18a-Kif18b chimera robustly targeted to microtubule ends, particularly to astral microtubules during mitosis (Fig 2C). These experiments demonstrate that the C terminus of Kif18b determines the localization of the motor domain to microtubule ends and that the Kif18a motor domain can become a Kif18b-like motor when fused to the C terminus of Kif18b.

**Figure 2:**
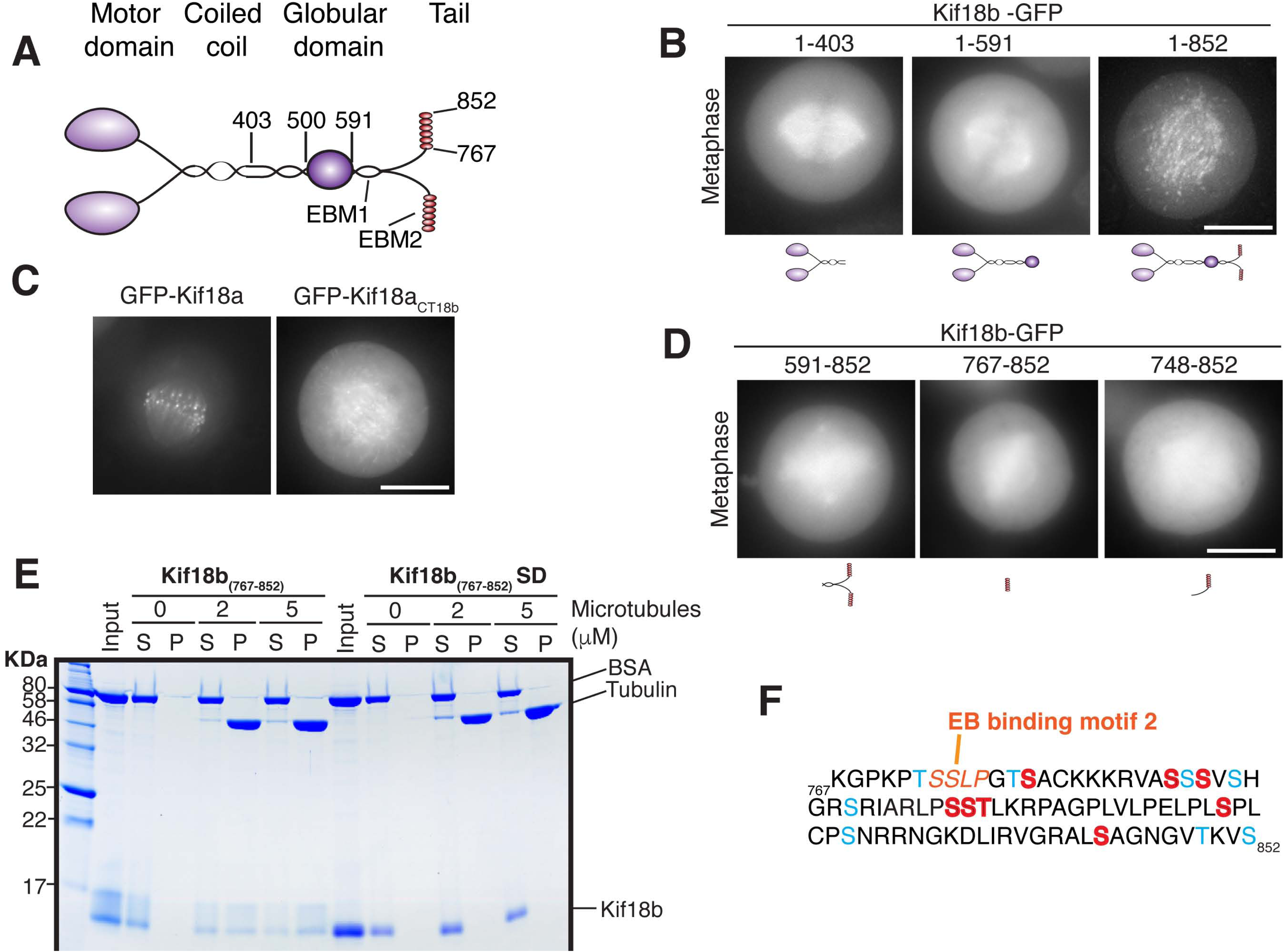
The C terminus of Kif18b is a microtubule-binding domain. (A) Schematic diagram representing Kif18b structural domain organization. <u>EB</u>-binding <u>m</u>otifs (EBM) are indicated. (B-D) Representative images of HeLa cells transfected with (B) full-length and truncated Kif18b motor (C) Kif18a and the chimeric Kif18a-Kif18b construct and (D) truncation of Kif18b C terminus. Scale bar 10 μm. (E) Coomassie-stained gel showing cosedimentation of the Kif18b C terminus, but not the phosphomimetic C terminus in the presence of 2 μM taxol-stabilized microtubules. (F) Amino acid sequence of the C terminus of Kif18b. EB-binding motif is highlighted in orange. The amino acids which are phosphorylated *in vivo* and mutated to aspartates in this study, and phosphorylatable residues are painted red and blue, respectively.

We next examined the role of the Kif18b C terminus Kif18b_591-852_ in Kif18b targeting. Surprisingly, Kif18b_591-852_ targeted to the spindle, rather than microtubule ends suggesting that this region contains a second microtubule-binding region. To test this, we generated a series of truncated constructs and found that the far C terminus, Kif18b_767-852_ is sufficient to localize to spindle microtubules (Fig 2D). In addition, we found that recombinant Kif18b_767-852_ binds to microtubules *in vitro* (Fig 2E). The C terminus of Kif18b is strongly phosphorylated in human cells during mitosis (Fig 2F) (Dephoure et al., 2008). To test whether phosphorylation regulates the microtubule binding properties of the Kif18b non-motor microtubule-binding domain, we generated a phosphomimetic mutant by substituting 8 residues within Kif18b_767-852_ that are phosphorylated *in vivo* to aspartates (Fig 2F)(Dephoure et al., 2008; Sharma et al., 2014). This phosphomimetic Kif18b_767-852_ domain did not bind to microtubules *in vitro* (Fig 2E), indicating that phosphorylation regulates the affinity of the Kif18b non-motor microtubule-binding domain for microtubules. Kif18b_767-852_ contains additional serines and threonines, which could be phosphorylated, although they have not been identified so far. Taken together, the C terminus of Kif18b provides a second microtubule-binding domain to Kif18b that is regulated by phosphorylation.

### The Kif18b C terminus is required for Kif18b accumulation at microtubule plus ends

Since Kif18b has a C-terminal non-motor microtubule-binding domain, we hypothesized it may regulate the microtubule-end targeting and functional properties of full-length Kif18b *in vivo*. To distinguish the contribution of the non-motor microtubule-binding domain and the EB-binding motifs to the plus end accumulation of Kif18b *in vivo*, we transfected Kif18b, Kif18bΔ_SXIP,_ mutating the last two residues in the putative EB-binding motifs to NN, and Kif18bΔ_C_ lacking the C-terminal microtubule-binding domain (767-852), into the Kif18b KO cells. Kif18b co-localized with EB1 at microtubule plus ends (Fig 3A-C). Kif18bΔ_SXIP_ had strongly reduced plus tip localization. However, we could still observe weak plus tip targeting (Fig 3A-C) as previously reported (Tanenbaum et al., 2011). Kif18bΔ_C_ was also strongly reduced at microtubule plus ends (Fig 3A-C), despite the presence of an upstream EB motif (Fig 2A). Importantly, Kif18b_SXIPmut2_, with mutations in just the C-terminal EB-binding motif (_773_SXIP_776_) still robustly localized to microtubule ends (Fig 2A, F, S3), indicating that the C-terminal microtubule-binding region and the EB-binding motifs play non-redundant targeting functions.

**Figure 3:**
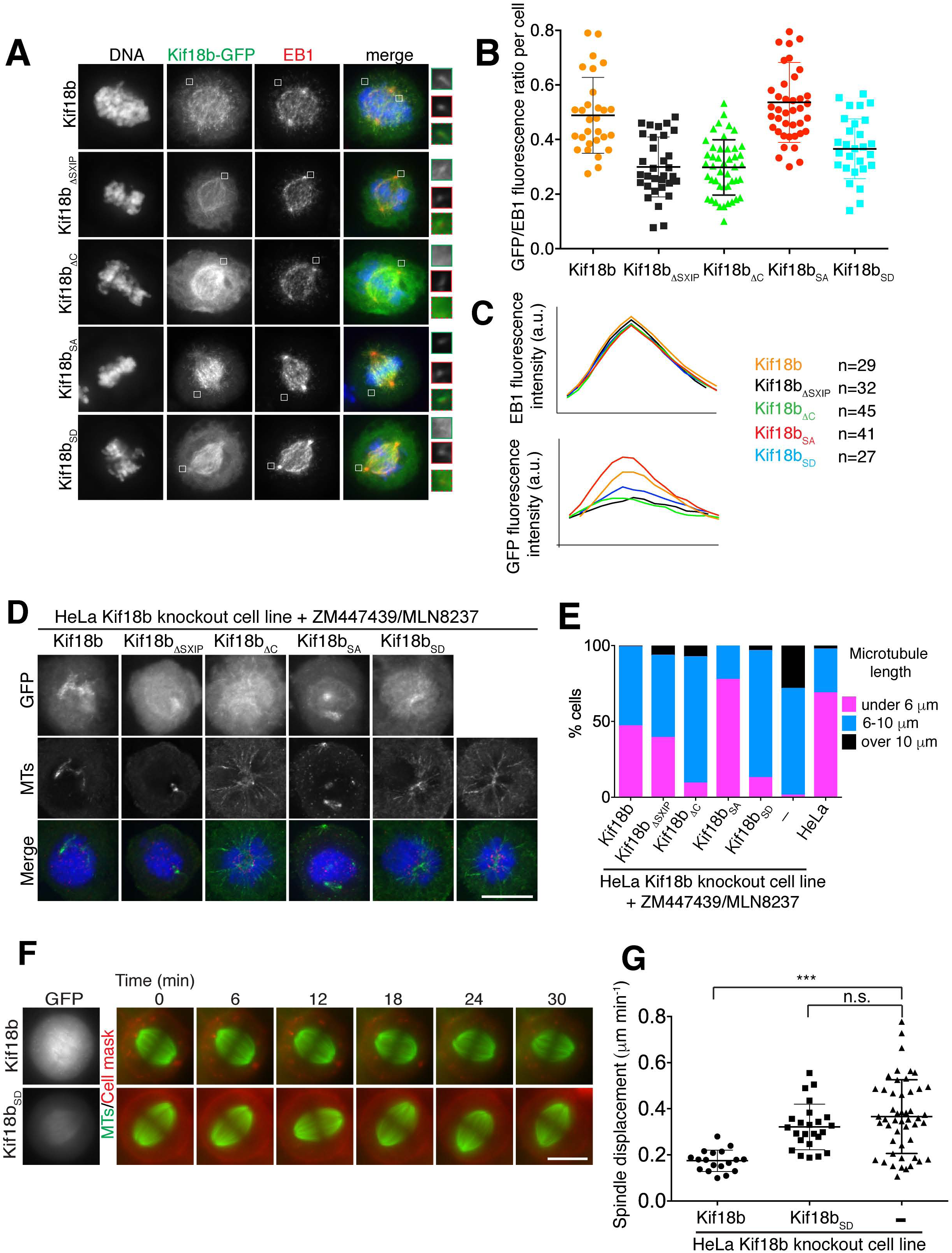
The C terminus of Kif18b contributes to Kif18b targeting to microtubule plus ends and microtubule length *in vivo*. (A) Representative immunofluorescence images of Kif18b-KO HeLa cells transfected Kif18b-GFP constructs and stained for EB1. Scale bar 10 μm. (B) Quantification of GFP/EB1 signal intensity ratio at microtubule plus end. Each data-point represents one cell for which on average 4 comet intensities have been measured and averaged (mean±sd). Kif18b-GFP transfection repeated twice, all others three times. (C) Averaged linescan profile of measured EB1 comets and their corresponding linescan profile for GFP-Kif18b constructs from cells measured in (B). (D) Representative immunofluorescence images of Kif18b-KO HeLa cells transfected with Kif18b constructs. Cells were treated with Aurora kinase A and B inhibitors, MLN8237 and ZM447439. (E) Quantification of microtubule length in cells treated with STLC and MLN8237/ZM447439 after transfection of Kif18b-GFP constructs (111>n>66). (F) Time-lapse imaging series of Kif18b-KO HeLa cells transfected with Kif18b-GFP and Kif18b_SD_-GFP, imaged with SiR-tubulin and CellMask dye. (G) Quantification of the displacement of the spindle from the center of the cell during metaphase. Each data point represents spindle displacement over 30-40 minutes. Error bars represent standard deviation. Asterix indicate Ordinary one-way Anova test significance value ****P<0.0001. Scale bars 10 μm.

Since mitotic phosphorylation decreases the affinity of the Kif18b non-motor microtubule-binding domain for microtubules, we hypothesized that phosphorylated Kif18b would accumulate less at microtubule plus ends, whereas a non-phosphorylatable Kif18b mutant would be enriched. To test this, we generated non-phosphorylatable (Kif18_SA_-GFP) and phosphomimetic (Kif18_SD_-GFP) mutants for the 8 previously identified phosphorylated sites (Dephoure et al., 2008). Kif18b_SA_ displayed increased targeting to microtubule plus tips. In contrast, Kif18b_SD_ showed severely reduced plus tip localization, indicating that mitotic phosphorylation of its C terminus controls its accumulation at microtubule ends (Fig 3A-C). Taken together, these results indicate that Kif18b utilizes both its C-terminal microtubule-binding domain and the EB-binding motifs to accumulate at microtubule plus ends. Phosphorylation of the C terminus of Kif18b reduces its plus-end accumulation, which could control its spatial distribution of Kif18b during mitosis.

### The C terminus of Kif18b is a requirement for spindle positioning

Next, we sought to test whether the C terminus of Kif18b was necessary for its ability to shorten microtubules and center the spindle in Kif18b KO cells. Kif18b is negatively regulated by Aurora kinases. Upon Aurora kinase small molecule inhibition, Kif18b depolymerizes microtubules (Tanenbaum et al., 2011). In our Hela Kif18b KO cell line, microtubules remain long in the presence of Aurora kinase inhibitor (Fig 1C, D). Kif18b and Kif18b_SA_ transfection restored the control of microtubule length, with shortened and depolymerized microtubules, as observed in HeLa cells (Fig 3D, E). In contrast, the ability of Kif18b_SD_ and Kif18bΔ_C_ to depolymerize microtubules was modest, suggesting the non-motor microtubule-binding domain of Kif18b plays a role in microtubule depolymerization. Kif18bΔ_SXIP_ displayed an intermediate ability to depolymerize microtubules, consistent with its weak localization to plus ends (Fig 3E). Finally, we tested whether the non-motor microtubule-binding domain of Kif18b was important for the ability of the kinesin to control spindle positioning. Kif18b restored correct spindle centering cells lacking Kif18b (Fig 3F, G). However, the ability of Kif18b_SD_ to center the spindle was strongly reduced, consistent with the fact that Kif18b_SD_ did not rescue proper microtubule length. These results indicate the presence of the Kif18b non-motor microtubule-binding domain is important for microtubule length regulation and spindle positioning. Phosphorylation of the second microtubule-binding domain of Kif18b reduces Kif18b accumulation at microtubule ends to prevent microtubule depolymerization.

### Kif18b is a processive plus end directed motor

The precise activity of Kif18b and how it reaches microtubule ends to shorten them remain unclear. To determine whether Kif18b acts as a *bona fide* microtubule depolymerase at microtubule ends, we expressed and purified Kif18b-GFP from insect cells (Fig S4A). We immobilized GMPCPP-stabilized microtubules on coverslips that had surface-adsorbed anti-tubulin antibodies at low density. Microtubules alone remained stable over time. In the presence of 50 nM MCAK, microtubules rapidly depolymerized at the rate 0.2 μm/min. In contrast, in the presence of Kif18b, the microtubules remained stable (Fig 4A, B). This indicates that Kif18b does not use the same mechanism as MCAK and Kip3 to depolymerize microtubules (Gardner et al., 2011; Su et al., 2011). To analyze the motile properties of Kif18b, we imaged Kif18b-GFP on stabilized microtubules using a single molecule TIRF microscopy-based assay. We observed Kif18b to move processively towards the plus ends of microtubules for long distances with an average speed of 349 ± 7 nm/s (mean +/-SE, Fig 4C, D). Kif18b could travel over distances longer than 7 μm (Fig S4B, C), with the run length limited by the length of the microtubule. As Kif18b-GFP reached the plus end of microtubules, it dwelled at the plus tip of microtubules for a significant amount of time (Fig 4E, F). Thus Kif18b recognizes the end of stabilized microtubules *in vitro* independently of EB proteins. These data indicate Kif18b is a fast and processive plus end-directed motor that targets and accumulates at the ends of microtubules *in vitro*.

**Figure 4:**
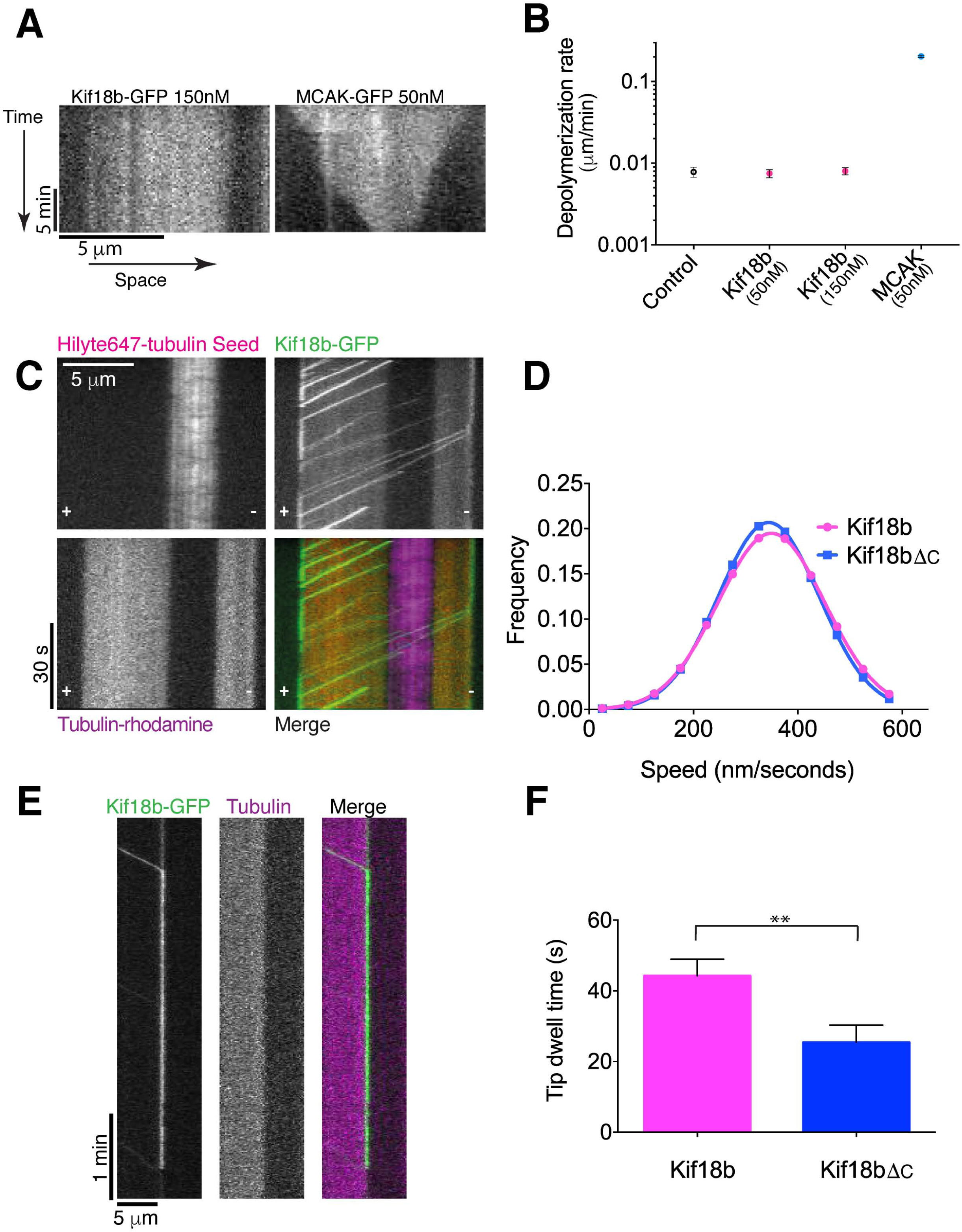
Kif18b is a fast processive plus end directed motor that uses its C terminus to accumulate at microtubule ends. (A, B) Kif18b is not a microtubule depolymerase. A) Example kymographs of GMP-CPP stabilised microtubules in the presence of 150 nM Kif18b and 50 nM MCAK. B) Depolymerisation rates for GMP-CPP stabilised microtubules with no added motor (n=47), 50 nM Kif18b (n=74), 150 nM Kif18b (n=50) and 50 nM MCAK (n=113) (mean±sem). (C) Kif18b is a plus end directed motor with a run length of >7µm. Polarity marked GMP-CPP stabilised microtubules (HiLyte and Rhodamine labelled) with Kif18b motors. Motors walk processively towards the plus end of the microtubules with motors pausing at the microtubules tips before dissociation. (D) The velocities of Kif18b (n=193) and Kif18bΔ_C_ (n=142) as measured from kymographs. (E) Example Kymograph showing Kif18b dwelling at a microtubule tip for over a minute. (F) Tip dwell time frequencies for Kif18b (n=103) and Kif18bΔ_C_ (n=73). Kif18bΔ_C_ dwells at microtubule tips for shorter times than full-length Kif18b (mean ±sem).

Given Kif18b accumulates at microtubule ends *in vivo* using its non-motor microtubule-binding region, we hypothesized the non-motor region may be important for this targeting property *in vitro*. To test the role of the non-motor microtubule-binding domain on the motility and microtubule-end targeting of full-length Kif18b, we purified Kif18bΔ_C_-GFP lacking the last 75 amino acids. Overall, the velocity of Kif18bΔ_C_ was similar to that of Kif18b (Fig 4C, D) and the motor remained highly processive (Fig S4). However, there was a remarkable reduction in its dwell time at the tips. More than half of Kif18bΔ_C_ delocalized from the plus end of microtubules within 8.5 s, in contrast to Kif18b (22.8 s) (Fig 4F). This indicates that the C-terminal non-motor microtubule-binding of Kif18b plays a role in the tethering of Kif18b, independently of EB1, once Kif18b reaches the microtubule end.

### Kif18b promotes microtubule catastrophe

Kif18b does not depolymerize stabilized microtubules unlike Kinesin-13 motors. To determine whether Kif18b regulates dynamic microtubules, we analyzed microtubule dynamic parameters and the length of dynamic microtubules in the presence and absence of Kif18b. First we observed that Kif18b walks processively to the ends of dynamic microtubules (Fig 5A). Kif18b then tracks the growing ends of microtubules. The microtubule growth rate and catastrophe rate increase with increasing amounts of Kif18b (Fig 5B, C). Inversely, the length of dynamic microtubule extensions decreased with increasing concentrations of Kif18b (Fig 5D, E). On average, extensions in the presence of tubulin alone were 1.7±0.1 μm long, while in the presence of 100nM Kif18b extensions were significantly shortened to 1.4±0.1 μm (mean+/-sem). Overall these results indicate that Kif18b reduces microtubule length by increasing the catastrophe rate of dynamic microtubules. In total, our data demonstrate that Kif18b uses its motile properties to promote remodeling of the cytoskeleton during mitosis, and positioning of the metaphase spindle. Kif18b is a processive plus end-directed motor, that uses its C-terminal microtubule-binding domain to accumulate at microtubule ends, where it promotes microtubule catastrophe and hence depolymerization. Mitotic phosphorylation of the C-terminal microtubule-binding domain provides a mechanism to spatially regulate the accumulation of Kif18b to microtubule ends, and to shorten sub-cellular populations of microtubules to ensure spindle positioning.

**Figure 5:**
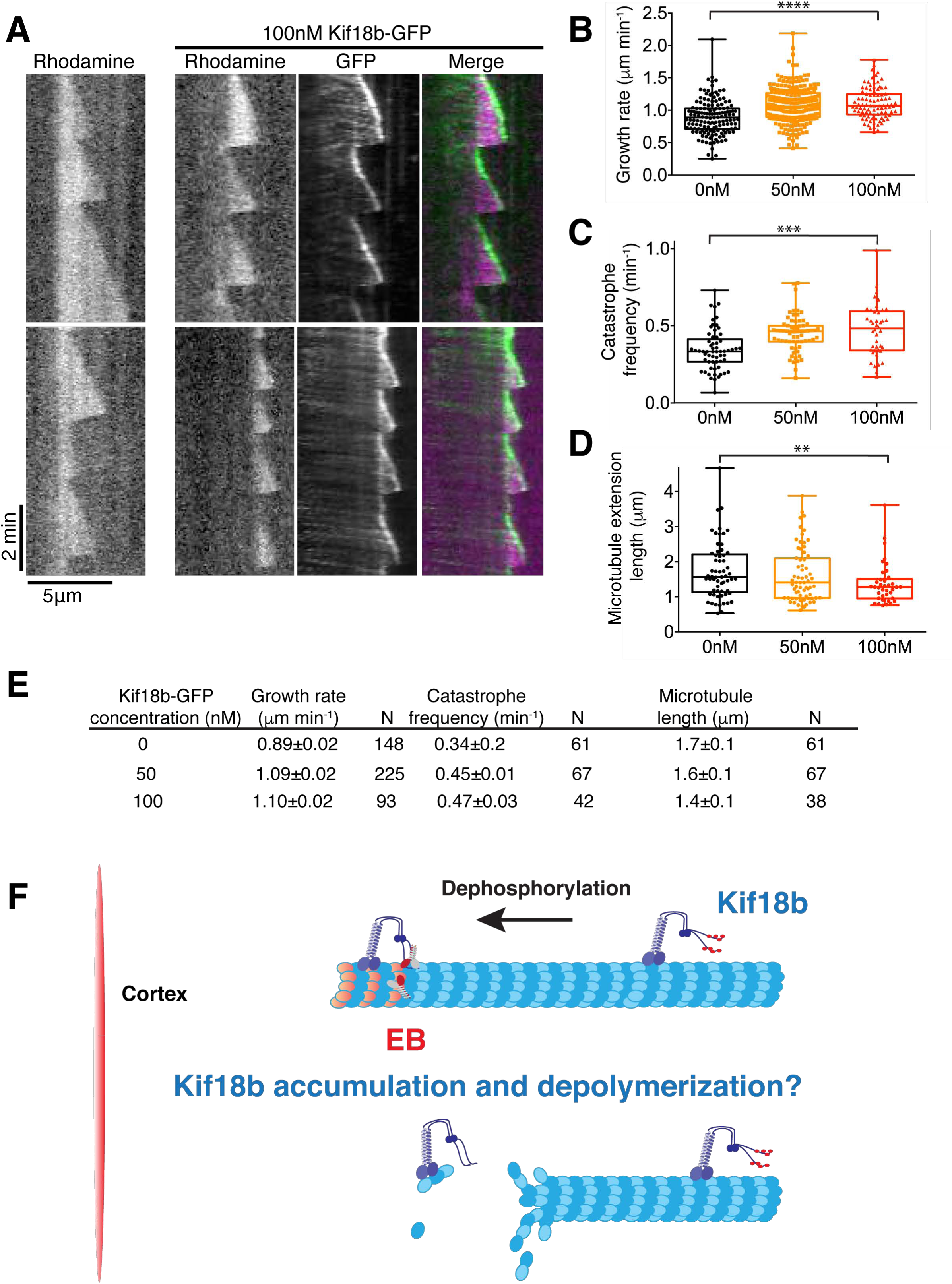
Kif18b accumulates at the tips of growing microtubules and acts as a catastrophe factor to reduce microtubule length. (A) Kif18b tracks the tips of growing microtubules. Example kymographs of rhodamine labeled dynamic microtubules in 7 µM tubulin with 0 nM and 100 nM dimeric Kif18b-GFP. (B) Measured growth rates of dynamic extensions in the presence of 7µM tubulin and 0, 50 and 100 nM dimeric Kif18b-GFP. Asterix indicate ANOVA-test significance value ****P<0.0001 (C) Catastrophe frequencies of dynamic microtubule extensions in the presence of 7 µM tubulin and 0, 50 and 100 nM dimeric Kif18b-GFP. Each data point corresponds to the catastrophe frequency for an individual microtubule. Asterix indicate KS test significance value ***P<0.001. (D) Average lengths of dynamic microtubule extensions in the presence of 7 µM tubulin and 0, 50 and 100 nM dimeric Kif18b-GFP. Each data point corresponds to the average length of the Rhodamine labeled extension from a Hilyte_647_ labeled seed over the course a kymograph. Asterix indicate KS test significance value **P<0.002 (B-D) Graphs are box and whiskers plots. Bars represent minimum and maximum points. (E) Table containing the microtubule dynamic parameters in the absence and presence of 50 and 100 nM Kif18b-GFP. (F) Schematic model of Kif18b walking towards the plus end of microtubules. Kif18b is phosphorylated close to centrosomes and chromosomes. As Kif18b reaches the cortex, the C terminus is dephosphorylated by cytoplasmic phosphatases, which allows Kif18b to accumulate at microtubule ends. Kif18b and MCAK could cooperate at microtubule plus ends to destabilize microtubules.

## Discussion

Microtubule depolymerization is an essential process in eukaryotes to ensure microtubules remodeling and facilitate microtubule-based processes. Kinesin-13 and Kinesin-8 are the major motor families responsible for stimulating microtubule depolymerization. While the mechanisms by which kinesin-13 motors destabilize microtubule ends is well established, the mechanism by which Kinesin-8 motors reduce microtubule length is more controversial and differs between members across species. Recent work proposed that Kif18b is a poorly processive motor which uses diffusion to reach microtubule ends, suggesting Kif18b tracks microtubule plus ends through a bind and release mechanism, similarly to other plus tip tracking proteins (Akhmanova and Steinmetz, 2008; Shin et al., 2015). We believe that a N-terminal GFP label may inhibit the motile properties of Kif18b, leading to its diffusive behavior (Shin et al., 2015). Indeed, our data show that Kif18b is a highly processive motor that walks to the plus ends of microtubules and negatively regulates dynamic microtubules by promoting catastrophe. The single molecule behavior of Kif18b, with high processivity and dwells at microtubule tips, is similar to the Kinesin-8s Kif18a and yeast Kip3, although Kif18b is faster (Stumpff et al., 2011; Su et al., 2011). However, its action at microtubule tips is distinct. Unlike Kif18a and Kip3, Kif18b does not appear to depolymerize stable microtubules or increase pausing (Mayr et al., 2007; Su et al., 2007; Locke et al., 2017). Additionally whilst Kif18a appears to dampen microtubule dynamics Kif18b achieves a reduction in microtubule length through an increase in microtubule dynamics, principally the catastrophe rate (Stumpff et al., 2011).

Kif18b uses a C-terminal non-motor microtubule-binding domain to recognize and accumulate at microtubule ends *in vitro*, similarly to multiple other kinesins, reviewed in (Welburn, 2013). These nucleotide-insensitive microtubule-binding regions play important roles in specifying the unique functions and targeting of kinesins. Interestingly, the C terminus of Kif18b does not have any effect on run length and processivity of the motor, instead affecting its microtubule tip targeting *in vivo* by reducing its ability to remain bound once it has reached microtubule tips. This domain is not conserved in sequence and appears to bind electrostatically to microtubules using its positively charged residues (pI=11.5). Given its long run length and its speed *in vitro*, Kif18b may walk along microtubules *in vivo*, until it reaches microtubule ends. Single molecule imaging of Kif18b *in vivo* measured that Kif18b moves processively on microtubules at a rate of 635 nm/s, which would allow it to reach the end of a polymerizing microtubule (Tanenbaum et al., 2014). We here show that *in vitro* Kif18b can track growing ends autonomously. *In vivo* Kif18b appears to use a dual plus end-targeting mechanism through microtubule end tethering and EB-binding, to robustly localize to the growing microtubule tip (Fig 5F). Endogenous Kif18b is enriched specifically at the ends of astral microtubules and is excluded from microtubules in the vicinity of chromosomes and centrosomes (Stout et al., 2011; Tanenbaum et al., 2011). Aurora kinases and Plk1 localize specifically to chromosomes and centrosomes during mitosis, thereby creating a spatial phosphorylation gradient for substrates (Liu et al., 2009; Ye et al., 2015). We propose that phosphorylation of the non-motor microtubule-binding domain of Kif18b by spatially-restricted mitotic kinases could exclude or at least reduce Kif18b from dwelling at the tips of microtubules in these regions. As Kif18b walks on microtubules away from the kinase activity gradient, it becomes dephosphorylated by cytoplasmic phosphatases. This dephosphorylation allows Kif18b to dwell at microtubule tips for longer, leading to subcellular accumulation of Kif18b at the plus tips of astral microtubules. In this manner, microtubule plus end localised Kif18b will be able to selectively destabilize these microtubules helping to ensure correct spindle positioning. (Fig 5F). *In vitro*, Kif18b_767-851_ is not an Aurora kinase substrate (data not shown). Future work should determine which mitotic kinases phosphorylate Kif18b (Dephoure et al., 2008).

Our study also sheds light on the mechanism by which Kif18b destabilizes microtubules. Our data suggest Kif18b does not destabilize microtubules to the extent of MCAK. However, Kif18b regulates microtubule plus end dynamics negatively by promoting catastrophe, resulting in a net reduction in the length of microtubules (Fig 5). Because Kif18b also modestly increases microtubule growth rate, microtubules are not completely depolymerized, unlike in the presence of MCAK. Rather microtubules have a tighter length distribution (Fig 5D). The processivity of Kif18b plays an important role ensuring the motor reaches microtubule ends to regulate them, similarly to Kip3 (Su et al., 2011; Varga et al., 2006; Varga et al., 2009). Kif18b is nuclear in interphase. Therefore its access to microtubules at mitotic onset and its spatial regulation contribute to the increase in microtubule dynamics and reduction of remodeling of the cytoskeleton in mitosis. In agreement with previous reports our data suggest MCAK and Kif18b may work in the same pathway to destabilize microtubules (Tanenbaum et al., 2011; Walczak et al., 2016). Future work should determine the mechanism by which Kif18b and MCAK cooperate using their distinct mechanisms to regulate microtubules during mitosis.

## Author contributions

AGK, TM and JW designed, performed experiments and analyzed data. JW wrote the manuscript.

## Acknowledgements

Plasmids pX330 and plasmid carrying mCherry-Cas9 were kind gifts of Feng Zhang and Peter Hohenstein. Bonsai Ndc80 antibody was a generous gift from Iain Cheeseman. J. W. is supported by a CRUK Career Development Fellowship (C40377/A12840), a FP7 Marie Curie Re-integration grant (33370) and a Royal Society grant (RG160003). We thank Sarah Young for help during the early stages of the project. We thank Thomas Surrey and Johanna Roostalu for help with preparation of coverslips for dynamic microtubule experiments and the Welburn lab for critical reading of the manuscript. J. W. is supported by a CRUK Career Development Fellowship (C40377/A12840). The Wellcome Trust Centre for Cell Biology is supported by core funding from the Wellcome Trust (203149).

---

**Supplementary figure 1:**
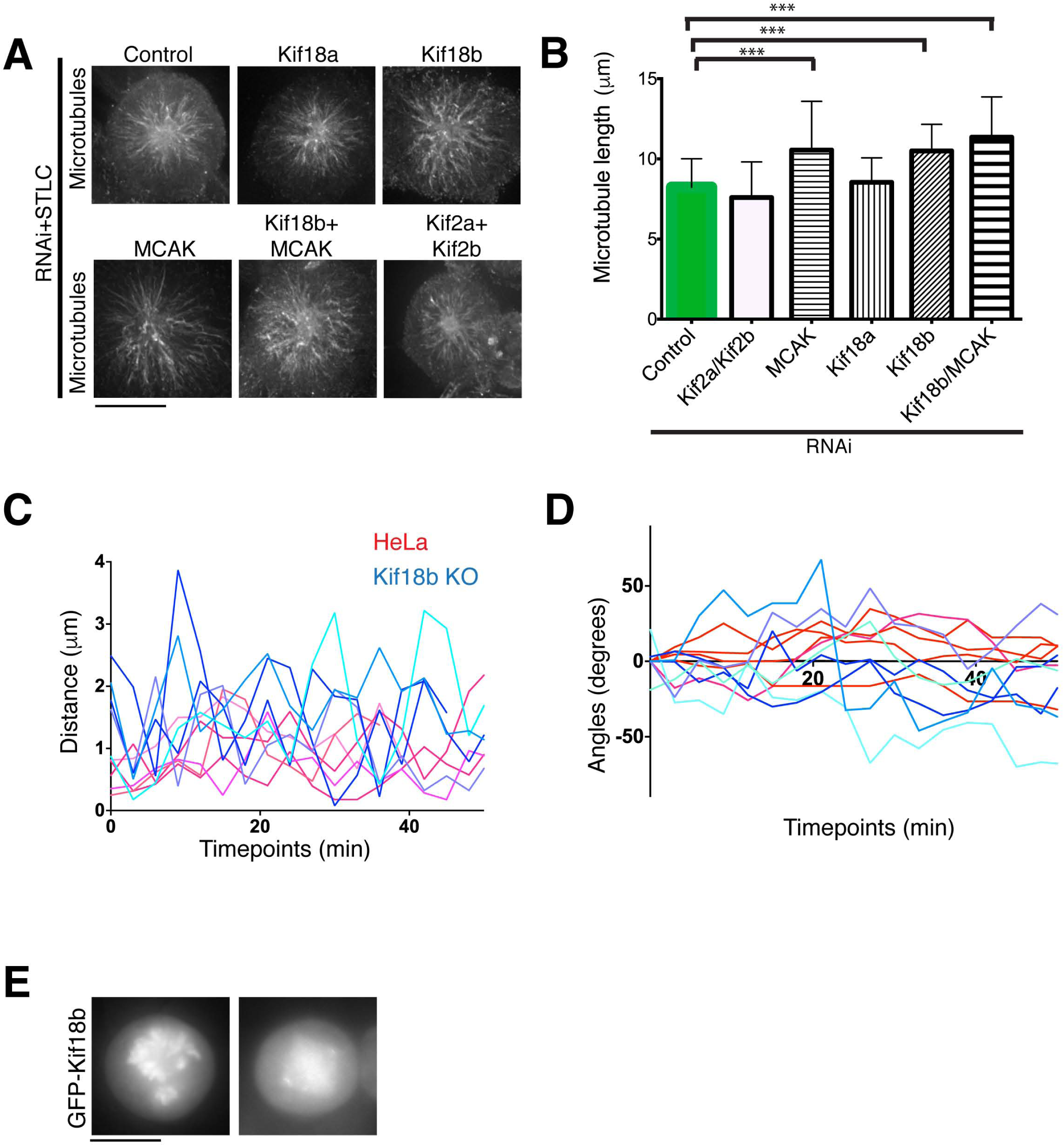
Effect of kinesin-8 and kinesin-13 motor family on microtubule length during mitosis. (A) Immunofluorescence images acquired using a tubulin antibody in U2OS cells depleted with control, kinesin-8 or kinesin-13 siRNA and treated with STLC. (B) Bar graph showing the average microtubule length for microtubules radiating from the centrosome (mean ± SD) for depletions in (A). Asterix indicate Ordinary one-way Anova test significance value ***P<0.001. Experiment performed quantified once. (C, D) Representative traces of individual spindles moving and rotating in the xy plane, with respect to the defined cell axis over time. (E) Representative images of transiently transfected GFP-Kif18b. Scale bars: 10 μm.

**Supplementary figure 2:**
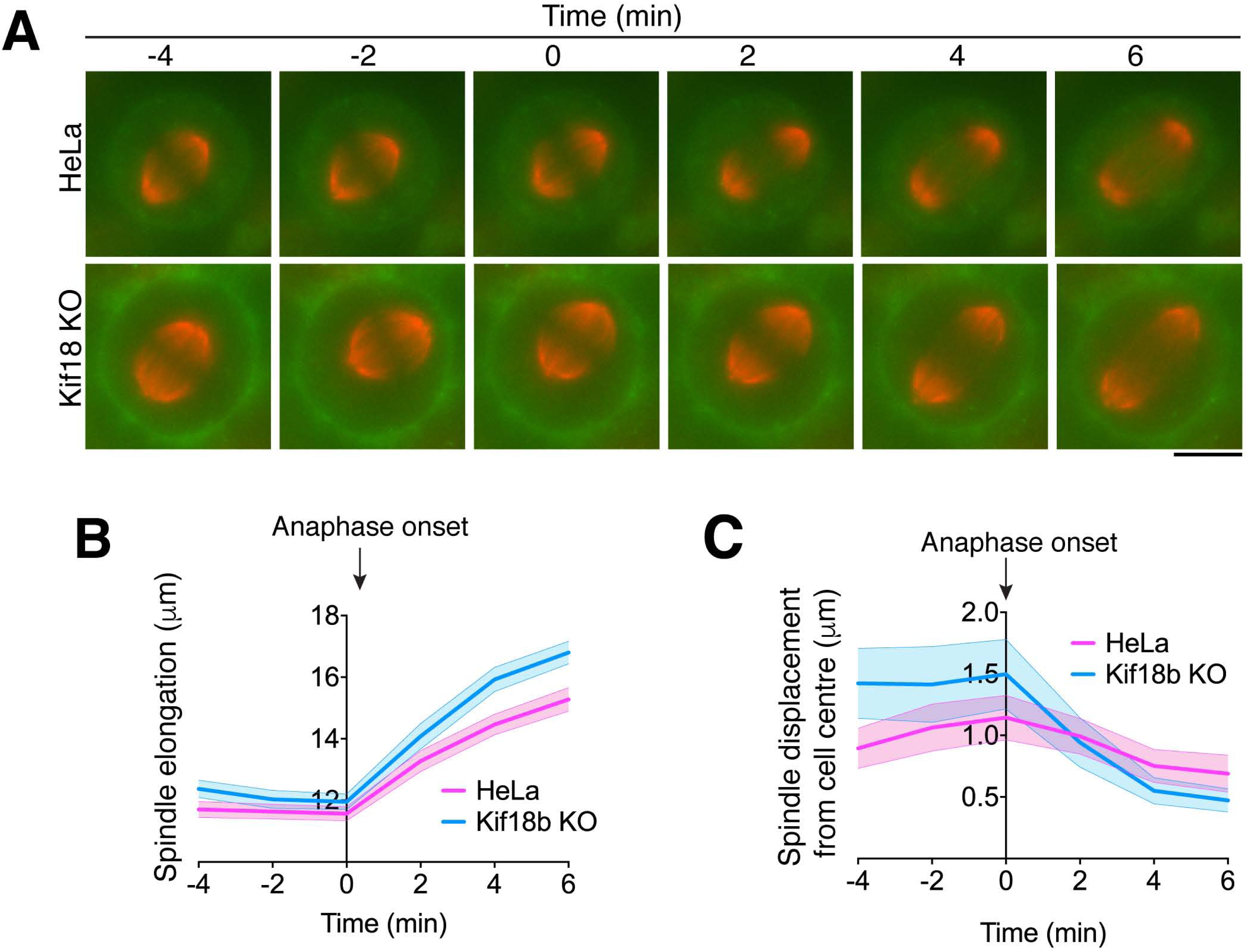
Spindle centering is corrected during anaphase. (A) Representative time-lapse images of cells going through mitosis incubated with SiR-tubulin (red) and CellMask dye (green). The zero timepoint is defined as the last frame before anaphase spindle elongation takes place. (B) Graph representing the average spindle length during the metaphase to anaphase transition and the corresponding SD for HeLa and Kif18b-KO HeLa cells (n= 64 and 44). (C) Graph showing the displacement of the centre of the spindle with respect of centre of the cell along the spindle elongation axis. Error bars represent 95% confidence intervals.

**Supplementary figure 3:**
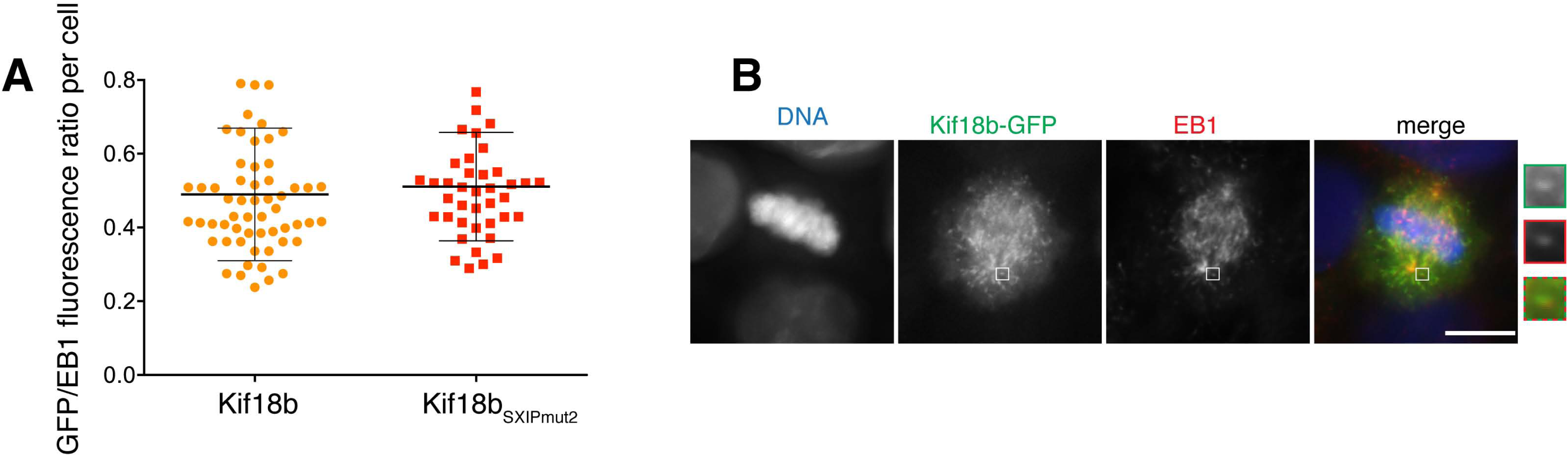
Disruption of EB-binding motif in Kif18b C terminus does not prevent Kif18b plus tip targeting. **(A)** Representative immunofluorescence images of Kif18b-KO HeLa cells transfected Kif18b_SXIPmut2_-GFP construct and stained for EB1. Scale bar 10 μm. (B) Quantification of GFP/EB1 signal intensity ratio at microtubule plus end for Kif18b-GFP and Kif18b_SXIPmut2_-GFP. Each data-point represents one cell for which at least 4 comet intensities have been measured and averaged.

**Supplementary figure 4:**
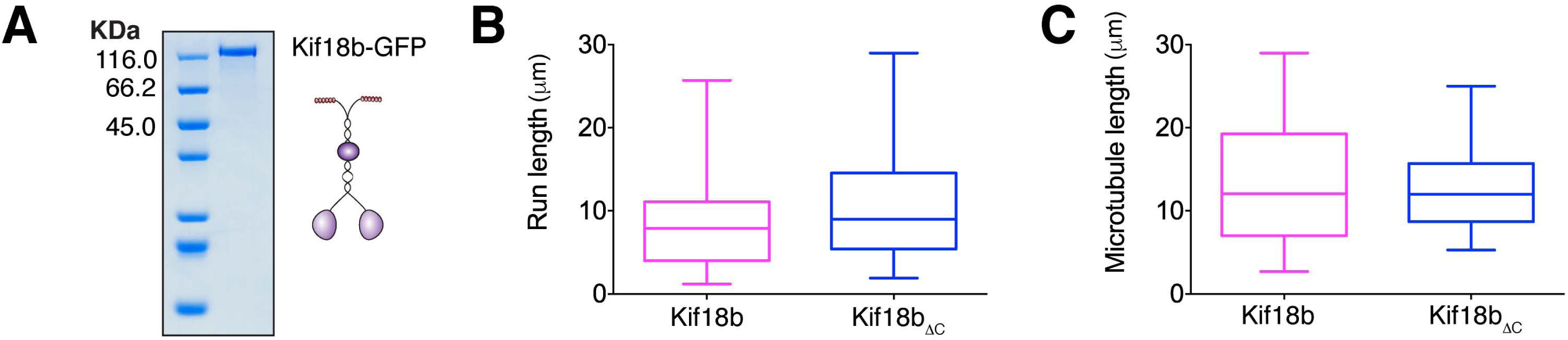
Kif18b has a long run length. (A) Coomassie-stained gel of insect cell-expressed and purified Kif18-GFP. (B, C) The run length of Kif18b (n=44, median = 7.9µm) and Kif18bΔ_C_ (n=33, median = 7.7µm) is limited by microtubule length with very few motors detaching from the microtubule before reaching the microtubule end. No significant difference can be seen in the run length between the two motors (B). The motor run lengths for the two motors were measured over similar microtubule populations (C).

**Supplementary Movie 1:** movie of a cell in metaphase displaying rocking and mispositioning of the spindle over time.

**Supplementary Movie 2:** Single molecule imaging of Kif18b-GFP walking towards the plus ends of microtubules. Microtubule seeds are labeled in blue (tubulin HiLyte_647_) and seeds were extended using rhodamine-tubulin to determine the polarity of microtubules.

## Methods

### Cloning

To assay the localization in cell culture of Kif18b subdomains, various constructs were generated from Kif18b transcript variant 1 (NM_001265577.1) and cloned into pBABE-puro containing an N or C-terminal LAP tag (Cheeseman and Desai, 2005). For constructs expressing a C terminal LAP-tag at the C terminus of Kif18b, an N-Terminal Kozak consensus sequence 5’ *ACCGTCGCCAC* 3’ was included upstream of Kif18b for enhanced translation initiation. The microtubule-binding domain of Kif18b was cloned into pET3aTr. The non-phosphorylatable (SA) and phosphomimetic (SD) mutants were synthesized as fragments (Mr Gene GmbH) and subcloned. Previously proposed EB-binding motifs were mutated using PCR amplification of the whole plasmid using complementary primers containing the mutations. Residues mutated are underlined: _653_SF<u>LP</u>_656, 773_SS<u>LP</u>_776, 799_AR<u>LP</u>_802_, although we do not consider this last sequence an EB-motif as A at this position is non-binding to EB (Buey et al, 2012). For recombinant expression in insect cells, full-length Kif18-GFP-His was cloned into pFL.

### Protein expression, purification and assays

Protein purification and microtubule co-sedimentation assay were performed as previously described (Talapatra et al., 2015). His-tagged proteins were purified using Ni-NTA–agarose beads (GE Healthcare Life Sciences) according to the manufacturer’s guidelines. Proteins were then purified using gel filtration chromatography pre-equilibrated in gel filtration buffer (For full-length Kif18b: 25 mM HEPES, pH 7.5, 150 mM NaCl, 300 mM KCl, 5 mM BME, 1 mM MgCl_2,_ 1 mM Na-EGTA, 1 mM ATP; for the Kif18b microtubule binding domain: 50 mM HEPES, pH 7.5, 150 mM NaCl, 5 mM BME). Analytical gel filtration chromatography was performed using either a Superdex 75 or a Superose 200 10/300 GL column (GE healthcare). For the Kif18b microtubule-binding domain, cleavage of the His tag was performed using the his-3C protease overnight at 4°C. For detection of Kif18b by Western blotting, an antibody against Kif18b_500-593_ was raised in rabbit (Covance) and affinity purified.

### Cell Culture and Transfection

HeLa cells were maintained in DMEM (Lonza) supplemented at 37°C in a humidified atmosphere with 5% CO_2_. Cells are monthly checked for mycoplasma contamination (MycoAlert detection kit, Lonza). For live imaging, cells were plated on 35 mm glass bottom microwell dishes (MatTek). Transient transfections were conducted using Effectene reagent (Qiagen) according to the manufacturer’s guidelines. RNAi experiments were conducted using RNAi MAX transfection reagent (Invitrogen) according to the manufacturer’s protocol. Cells visualized after 24-48h. Previously published siRNAs were used to deplete Kif18b (Kif18b on-Target plus SMARTpool siRNA Dharmacon)(Tanenbaum et al., 2011).

### CRISPR Cas9 knockout

The Kif18b guide RNA *CACCC*ACGCTGCAAGTAGTGGTAC was designed against the Kif18b 5’ exon using http://crispr.mit.edu. We transfected mCherry-*Streptococcus pyogenes* Cas9 and the plasmid pX330 containing a single guide RNA targeting the first exon of the Kif18b gene in HeLa cells. Double stranded DNA breaks are generated in the targeted exon such that repairs of these cuts can generate indels that disrupt the open reading frame and abolish protein synthesis (Cong et al., 2013). We then selected the monoclonal cell lines that lacked Kif18b and verified the depletion was specific to Kif18b. After 48 hours, single cells were sorted into 96 well-plates. Clonal cell lines were screened by Western blot for Kif18b depletion.

### Microscopy

For live-cell imaging, HeLa cells were transferred to Leibowitz L15 media (Life Technologies) supplemented with 10% FBS + penicillin/streptomycin (Gibco) and imaged in a 37°C using a DeltaVision core microscope (Applied Precision) equipped with a CoolSnap HQ2 CCD camera or with an EMCCD camera. For live-cell imaging, 4-10 z-sections were acquired at 0.5-1 µm steps using a 60x or 100x objective lens. For fixed imaging, 10-20 z-sections were acquired at 0.2-0.5 µm. The cells were imaged every 3 min. The spindle elongation assay was performed as previously described (Young et al., 2014). To visualize microtubules and the cell cortex, 20-100nM SiR dye (SpiroChrome) and 1:2000-1:40000 dilution of Cell Mask Orange (Thermo Fisher Scientific) were used for 3 h and 5 min, respectively. For immunofluorescence, cells were then washed in PBS and fixed by one of two methods, either fixed in methanol for 10 minute at −20°C and then permeabilised with cooled acetone for 1 minute at –20°C, or fixed in 3.8% formaldehyde in PHEM buffer (60 nM Pipes, 25 mM HEPES, 10 mM EGTA, 2 mM MgSO_4_, pH 7.0) for 20 minutes. Immunofluorescence in human cells was conducted using antibodies against mouse anti-β-tubulin (Sigma, 1:1000), mouse EB1 (BD transduction laboratories, 1:400) and rabbit Ndc80 antibodies (1:1000). For experiments with small molecule inhibitors, the proteasome inhibitor MG132, ZM447439, MLN8237 and the Eg5 inhibitor STLC were used at a final concentration of 10 μM, 2 μM, 300 nM and 5 μM respectively. Only cells expressing low levels of Kif18b were analyzed to avoid artifacts due to overexpression. For microtubule length measurement and GFP signal visualization 20 z-sections and 3 z-sections were acquired at 0.2-0.3 µm and 1 µm step size, respectively.

### TIRF microscopy

For experiments using stablised microtubule seeds silanised coverslips were prepared as in (Bechstedt et al., 2011) except that in place of treatment with Piranha solution coverslips were incubated overnight in 12% hydrochloric acid at 50°C. For experiments using dynamic microtubules, coverslips were prepared with a with a PEG-Biotin surface (Bieling et al., 2010). Flow chambers were formed using double sided sticky tape, a silanised coverslip and a microscopy slide. Flow chambers were 7-8 µl in volume.

All experiments were carried out in BRB80 buffer (80 mM K.Pipes, 1 mM MgCl_2_, 1 mM EGTA, pH6.8). For single molecule assays anti-beta tubulin antibodies (T7816, Sigma Aldrich) at a 1:10 dilution were first introduced to the chamber. The surface was then blocked with 1% pluronic F-127 (Sigma Aldrich) and GMP-CPP (Jena Bioscience) stabilized 7% Rhodamine-labeled microtubules (Cytoskeleton) were then bound to the glass surface via the antibodies. The surface was then further blocked with 1 mg/ml casein (C7906, Sigma Aldrich). Assay buffer consisted of BRB80 with 1 mM ATP, 0.5 mg/ml casein and an oxygen scavenging system (0.2 mg/ml glucose oxidase, 0.035 mg/ml catalase, 4.5 mg/ml glucose, 140 mM β-mercaptoethanol). 2-10 nM of dimeric Kif18b-GFP-His or Kif18bΔ_C_-GFP-His in assay buffer was introduced to the flow chamber, the chamber was then sealed with nail varnish and imaged immediately at 37°C.

For dynamic microtubule experiments the surface was first blocked with 5% pluronic F-127 before introduction of 50 µg/ml NeutrAvidin (Invitrogen). The chamber was then washed with 1 mg/ml casein and 1% Tween-20 (SLS) before introduction of GMP-CPP stabilised microtubule seeds with 7% Hilyte_647_ label (Cytoskeleton) and 7% Biotin label (Cytoskeleton). Microtubule seeds were allowed to bind to the surface before further blocking with 1 mg/ml casein and 1% Tween-20. Final assay buffer consisted of BRB80 with 1 mM ATP, 1mM GTP, 0.5 mg/ml casein, 0.5% Tween-20, 7μM tubulin (6% rhodamine-label) and an oxygen scavenging system. Up to 100nM of dimeric Kif18b-GFP-His in assay buffer was introduced to the flow chamber, the chamber was then sealed with nail varnish and imaged at 30°C.

Imaging was performed on a Zeiss Axio Observer Z1 TIRF microscope using a 100x NA1.46 objective and a Photometrics Evolve Delta EMCCD camera controlled by Zeiss Zen Blue software. Single molecule imaging was performed for up to 5 minutes at 2 fps. For calculation of tip dwell times a frame rate of 1 fps was used. Depolymerisation assays were performed over 15 minutes at 4 fpm. Microtubule dynamics were imaged at 0.25 fps for up to 15 minutes. Kymographs were produced using Image J and run lengths and velocities of motors and microtubule growth rates were measured from these kymographs. Tip dwell times were calculated from 2-colour kymographs where a motor could be seen to reach an unoccupied microtubule tip. Catastrophe frequencies were calculated from the length of and number of catastrophes seen in individual kymographs. Microtubule extension lengths are averages over individual kymographs. Normality was checked using an Anderson-Darling test, p<0.05 was used to reject the null hypothesis. A Kolmogorov-Smirnov test was used to compare data that was not normally distributed (Fig 5C, D). Images were stored and vizualised using an OMERO.insight client (OME).

### Statistics and reproducibility

Statistical analyses were performed using GraphPad Prism 6.0 or R software. No statistical method was used to predetermine sample size. No samples were excluded from the analyses. The investigators were not blinded to allocation during experiments and outcome assessment. All experiments were performed and quantified from at least three independent experiments, unless specified and the representative data are shown.

### Data availability

All data supporting the findings of this study are available from the corresponding author on request.

## References

Akhmanova, A., and M.O. Steinmetz. 2008. Tracking the ends: a dynamic protein network controls the fate of microtubule tips. Nat Rev Mol Cell Biol. 9:309–322.

Bakhoum, S.F., S.L. Thompson, A.L. Manning, and D.A. Compton. 2009. Genome stability is ensured by temporal control of kinetochore-microtubule dynamics. Nat Cell Biol. 11:27–35.

Bechstedt, S., M. Wieczorek, M. Noujaim, and G.J. Brouhard. 2011. Variations on the single-molecule assay for microtubule-associated proteins and kinesins. Methods Mol Biol. 777:167–176.

Belmont, L.D., A.A. Hyman, K.E. Sawin, and T.J. Mitchison. 1990. Real-time visualization of cell cycle-dependent changes in microtubule dynamics in cytoplasmic extracts. Cell. 62:579–589.

Bieling, P., I.A. Telley, C. Hentrich, J. Piehler, and T. Surrey. 2010. Fluorescence microscopy assays on chemically functionalized surfaces for quantitative imaging of microtubule, motor, and +TIP dynamics. Methods Cell Biol. 95:555–580.

Buey, R.M., I. Sen, O. Kortt, R. Mohan, D. Gfeller, D. Veprintsev, I. Kretzschmar, J. Scheuermann, D. Neri, V. Zoete, O. Michielin, J.M. de Pereda, A. Akhmanova, R. Volkmer, and M.O. Steinmetz. 2012. Sequence determinants of a microtubule tip localization signal (MtLS). J Biol Chem. 287:28227–28242.

Cheeseman, I.M., and A. Desai. 2005. A combined approach for the localization and tandem affinity purification of protein complexes from metazoans. Sci STKE. 2005:pl1.

Cong, L., F.A. Ran, D. Cox, S. Lin, R. Barretto, N. Habib, P.D. Hsu, X. Wu, W. Jiang, L.A. Marraffini, and F. Zhang. 2013. Multiplex genome engineering using CRISPR/Cas systems. Science. 339:819–823.

Dephoure, N., C. Zhou, J. Villen, S.A. Beausoleil, C.E. Bakalarski, S.J. Elledge, and S.P. Gygi. 2008. A quantitative atlas of mitotic phosphorylation. Proc Natl Acad Sci U S A. 105:10762–10767.

Gardner, M.K., M. Zanic, C. Gell, V. Bormuth, and J. Howard. 2011. Depolymerizing kinesins Kip3 and MCAK shape cellular microtubule architecture by differential control of catastrophe. Cell. 147:1092–1103.

Garzon-Coral, C., H.A. Fantana, and J. Howard. 2016. A force-generating machinery maintains the spindle at the cell center during mitosis. Science. 352:1124–1127.

Gupta, M.L., Jr., P. Carvalho, D.M. Roof, and D. Pellman. 2006. Plus end-specific depolymerase activity of Kip3, a kinesin-8 protein, explains its role in positioning the yeast mitotic spindle. Nat Cell Biol. 8:913–923.

Kiyomitsu, T., and I.M. Cheeseman. 2012. Chromosome- and spindle-pole-derived signals generate an intrinsic code for spindle position and orientation. Nat Cell Biol. 14:311–317.

Kiyomitsu, T., and I.M. Cheeseman. 2013. Cortical dynein and asymmetric membrane elongation coordinately position the spindle in anaphase. Cell. 154:391–402.

Liu, D., G. Vader, M.J. Vromans, M.A. Lampson, and S.M. Lens. 2009. Sensing Chromosome Bi-Orientation by Spatial Separation of Aurora B Kinase from Kinetochore Substrates. Science.

Locke, J., A.P. Joseph, A. Pena, M.M. Mockel, T.U. Mayer, M. Topf, and C.A. Moores. 2017. Structural basis of human kinesin-8 function and inhibition. Proc Natl Acad Sci U S A. 114:E9539–E9548.

Manning, A.L., N.J. Ganem, S.F. Bakhoum, M. Wagenbach, L. Wordeman, and D.A. Compton. 2007. The kinesin-13 proteins Kif2a, Kif2b, and Kif2c/MCAK have distinct roles during mitosis in human cells. Mol Biol Cell. 18:2970–2979.

Mayr, M.I., S. Hummer, J. Bormann, T. Gruner, S. Adio, G. Woehlke, and T.U. Mayer. 2007. The human kinesin Kif18A is a motile microtubule depolymerase essential for chromosome congression. Curr Biol. 17:488–498.

Rusan, N.M., C.J. Fagerstrom, A.M. Yvon, and P. Wadsworth. 2001. Cell cycledependent changes in microtubule dynamics in living cells expressing green fluorescent protein-alpha tubulin. Mol Biol Cell. 12:971–980.

Samora, C.P., B. Mogessie, L. Conway, J.L. Ross, A. Straube, and A.D. McAinsh. 2011. MAP4 and CLASP1 operate as a safety mechanism to maintain a stable spindle position in mitosis. Nat Cell Biol. 13:1040–1050.

Savoian, M.S., and D.M. Glover. 2010. Drosophila Klp67A binds prophase kinetochores to subsequently regulate congression and spindle length. J Cell Sci. 123:767–776.

Sharma, K., R.C. D’Souza, S. Tyanova, C. Schaab, J.R. Wisniewski, J. Cox, and M. Mann. 2014. Ultradeep human phosphoproteome reveals a distinct regulatory nature of Tyr and Ser/Thr-based signaling. Cell Rep. 8:1583–1594.

Shin, Y., Y. Du, S.E. Collier, M.D. Ohi, M.J. Lang, and R. Ohi. 2015. Biased Brownian motion as a mechanism to facilitate nanometerscale exploration of the microtubule plus end by a kinesin-8. Proc Natl Acad Sci U S A. 112:E3826–3835.

Siller, K.H., and C.Q. Doe. 2009. Spindle orientation during asymmetric cell division. Nat Cell Biol. 11:365–374.

Stout, J.R., A.L. Yount, J.A. Powers, C. Leblanc, S.C. Ems-McClung, and C.E. Walczak. 2011. Kif18B interacts with EB1 and controls astral microtubule length during mitosis. Mol Biol Cell. 22:3070–3080.

Stumpff, J., Y. Du, C.A. English, Z. Maliga, M. Wagenbach, C.L. Asbury, L. Wordeman, and R. Ohi. 2011. A Tethering Mechanism Controls the Processivity and Kinetochore-Microtubule Plus-End Enrichment of the Kinesin-8 Kif18A. Mol Cell. 43:764–775.

Stumpff, J., G. von Dassow, M. Wagenbach, C. Asbury, and L. Wordeman. 2008. The kinesin-8 motor Kif18A suppresses kinetochore movements to control mitotic chromosome alignment. Dev Cell. 14:252–262.

Su, X., W. Qiu, M.L. Gupta, Jr., J.B. Pereira-Leal, S.L. Reck-Peterson, and D. Pellman. 2011. Mechanisms underlying the dual-mode regulation of microtubule dynamics by kip3/kinesin-8. Mol Cell. 43:751–763.

Talapatra, S.K., B. Harker, and J.P. Welburn. 2015. The C-terminal region of the motor protein MCAK controls its structure and activity through a conformational switch. Elife. 4.

Tanenbaum, M.E., L.A. Gilbert, L.S. Qi, J.S. Weissman, and R.D. Vale. 2014. A proteintagging system for signal amplification in gene expression and fluorescence imaging. Cell. 159:635–646.

Tanenbaum, M.E., L. Macurek, A. Janssen, E.F. Geers, M. Alvarez-Fernandez, and R.H. Medema. 2009. Kif15 cooperates with eg5 to promote bipolar spindle assembly. Curr Biol. 19:1703–1711.

Tanenbaum, M.E., L. Macurek, B. van der Vaart, M. Galli, A. Akhmanova, and R.H. Medema. 2011. A Complex of Kif18b and MCAK Promotes Microtubule Depolymerization and Is Negatively Regulated by Aurora Kinases. Curr Biol. 21:1356–1365.

Varga, V., J. Helenius, K. Tanaka, A.A. Hyman, T.U. Tanaka, and J. Howard. 2006. Yeast kinesin-8 depolymerizes microtubules in a length-dependent manner. Nat Cell Biol. 8:957–962.

Varga, V., C. Leduc, V. Bormuth, S. Diez, and J. Howard. 2009. Kinesin-8 motors act cooperatively to mediate length-dependent microtubule depolymerization. Cell. 138:1174–1183.

Walczak, C.E., H. Zong, S. Jain, and J.R. Stout. 2016. Spatial regulation of astral microtubule dynamics by Kif18B in PtK cells. Mol Biol Cell. 27:3021–3030.

Welburn, J.P. 2013. The molecular basis for kinesin functional specificity during mitosis. Cytoskeleton (Hoboken). 70:476–493.

Welburn, J.P., and I.M. Cheeseman. 2012. The microtubule-binding protein Cep170 promotes the targeting of the kinesin-13 depolymerase Kif2b to the mitotic spindle. Mol Biol Cell. 23:4786–4795.

Ye, A.A., J. Deretic, C.M. Hoel, A.W. Hinman, D. Cimini, J.P. Welburn, and T.J. Maresca. 2015. Aurora A Kinase Contributes to a Pole-Based Error Correction Pathway. Curr Biol. 25:1842–1851.

Young, S., S. Besson, and J.P. Welburn. 2014. Length-dependent anisotropic scaling of spindle shape. Biology open.

